# Quantifying Costs of Urbanisation: Wetland Loss and Impacts in a Rapidly Developing Global City

**DOI:** 10.1101/2020.06.22.127365

**Authors:** Harriet Gabites, Ricky-John Spencer

## Abstract

1. As cities grow, natural ecosystems decline through conversion to urban environments. Cities are often viewed as biodiversity wastelands, but they can be hotspots of global biodiversity. Urban biodiversity emphasises two fundamentals. First, people living in cities enjoy wildlife and second, there is virtually no planning for species that co-inhabit our cities. If urban biodiversity was a significant part of planning, then we would be far better at integrating green infrastructure into expanding urban environments.
2. Wetlands are among the most important and productive ecosystems in the world. They are the main suppliers of fresh water for human use and provide habitat to critical fauna and flora. In urban areas they are a vital link to nature and social cohesion. Currently, there is an absence of wetland inventory quantifying loss and changes overtime. Hence the broad impacts of urbanisation on wetland loss are difficult to assess.
3. We explored wetland loss and created a wetland inventory for Western Sydney, Australia, one of the world’s fastest growing urban regions. We used satellite imagery to determine wetland number and type, and calculated changes in wetland surface area from 2010-2017. Broad changes to land use were also quantified. We developed species distribution models of a common urban wetland turtle (*Chelodina longicollis*) that people interact with regularly or have as pets. *Chelodina longicollis* utilises both aquatic and terrestrial environments, and we determined if changes in distribution were associated with changes in the wetland inventory and urbanisation.
4. Most local government areas (LGA) experienced a decrease in wetland surface area from 2010-2017, ranging from -1% (Cumberland) to -21% (Blacktown). Majority of LGAs experienced a decrease in wetland density, with wetland densities declining by 25% (Blacktown). All LGAs experienced an increase in urban land use, ranging from 3-12%, which was associated with high rates of wetland loss.
5. Changes in turtle distribution over the decade reflects a southern distribution shift away from where wetland losses were concentrated. We estimated that ∼40,000 individual turtles were displaced or killed due to wetland loss and urbanisation.
6. Urbanisation was the leading cause of wetland loss and degradation in Western Sydney between 2010 and 2017. Wetlands provide critical green infrastructure and significant green space for social cohesion in urban areas. Integration of current wetlands, or the creation of functional wetlands, is key for sustainable development of urban landscapes. Urban wetlands (natural and constructed) may provide “biodiversity arks” for endangered species and facilitate community led conservation programs.

## Introduction

The post-COVID19 built environment must avoid a long-term legacy where people fear public places, such as cities, workplaces, and other people. Hence urban design approaches known as tactical urbanism are a way to create or reconfigure public spaces that are both safe and social, as well as, enhance urban ecology and biodiversity. Similarly, government policies, such as the Natural Environment White Paper (NEWP) and the National Planning Policy Framework in the UK recognise the essential contribution and services that our natural environment can provide in the move towards more sustainable development. NEWP recognises that the natural environment underpins economic prosperity, health and wellbeing and government policies throughout the world are also recognising the importance of Green Infrastructure in the Water-Energy-Food nexus. As cities grow, natural ecosystems decline through their conversion to urban environments. Cities are often viewed as biodiversity wastelands, but they can be hotspots of global biodiversity. Cities represent opportunities for forwarding global biodiversity and sustainability goals and urban biodiversity emphasizes two fundamental truisms (Shaffer 2018). First, people living in cities enjoy wildlife and second, there is virtually no planning or foresight guiding the species composition that co-inhabit our cities (Shaffer 2018). If it was a significant part of planning, then we could further integrate green infrastructure into expanding urban environments.

Most urbanisation is occurring in regions identified as biodiversity hotspots (Seto et al. 2012), and is significantly impacting on ecological functioning because of habitat destruction, degradation, and fragmentation; increased levels of pollution; and alterations of natural disturbance regimes and ecosystem processes, such as water and nutrient cycling (Luck 2007, Grimm et al. 2008, Nilon et al. 2017). As a result, the abundance of flora and fauna is substantially reduced in urban areas compared with that in nonurban habitats (Aronson et al. 2014). Reductions in urban biodiversity have consequences for human well-being, reducing the benefits people obtain from nature at individual and community levels (Brown and Grant 2005, Nilon et al. 2017). However, cities can support significant levels of biodiversity, including endangered species, and can play an important role in conservation (Aronson et al. 2014, Ives et al. 2015).

A significant component of Green Infrastructure are urban wetlands (Ling *et al*. 2018). Wetlands play a vital role in ecosystem services; specifically, through provisioning, regulating, cultural and supporting services (McInnes 2013), including the supply of food and fresh water, groundwater discharge, pollution control, and climate regulation, but also cultural services including spiritual and scientific/educational benefits (Millennium Ecosystem Assessment 2005). Wetlands support 10% of the world’s biodiversity, despite representing only 0.01% of the world’s habitats (Balian *et al*. 2008, McInnes 2013) and are the most threatened ecosystems (Balmford *et al*. 2002; Zhang *et al*. 2010). Thirty-two percent of the world’s amphibian species are listed as threatened and wetlands are essential breeding habitat. Waterbirds depend on the availability of quality wetlands and 12% are now considered threatened, with habitat loss considered a significant threat (Houlahan *et al*. 2000; Stuart *et al*. 2004). However, we are only scratching at the surface assessing overall biodiversity loss because of reductions in wetland quantity and quality as a result of urbanisation (Hull 2016). Wetland loss is occurring at a faster rate than wetland reserves are created (McCauley, Jenkins & Quintana-Ascencio 2013) and urban wetlands provide critical habitat for some of the worlds most endangered species. However, constructed urban wetland reserves are often in areas where wetland connectivity is limited, and they are designed solely for recreational uses or to alleviate risks of flooding (Hull 2016). The true extent of wetland loss is unknown in most countries because comprehensive national inventories rarely exist. Globally, intermittent estimates of wetland loss exist but due to lack of standardisation and consistent definition of the term wetland, the estimates cannot be directly compared. South America, Russia and Africa record no wetland data (Hu *et al*. 2017). Some European countries have produced sporadic inventories but estimates would not be comparable to USA estimates which have been steadily logged since 1979 through a National Wetland Inventory (NWI) (Ling *et al*. 2018; Hull 2016). Wetland loss in Australia is unknown because a comprehensive national inventory does not exist, however a NSW state-wide inventory was in produced in 2004, but its focus was on the larger inland wetlands (Ling *et al*. 2018).

Urban wildlife has positive impacts on cities and urban residents by providing ecosystem services and inspire a feeling of connection with nature. Freshwater turtles, such as the eastern long-necked turtle (*Chelodina longicollis*), are common in urban wetlands and are regarded as indicators of water and ecosystem quality (Blamires & Spencer 2013). They play a critical role in ecosystem function through energy flow and nutrient cycling by consuming vast quantities of carrion. Historically, they were widespread throughout eastern Australia in large biomasses (Thompson 1993). They inhabit almost all types of urban wetlands (DPIE 2018), but populations are declining due to predation from invasive species and threats from urbanisation, such as road mortality from vehicle impacts (Santori *et al*. 2018). Long-necked turtles regularly move large distances over land to new wetlands for nesting and foraging (Thompson 1993), hence removal of wetlands increases the distances that turtles are required to move between wetlands. This nomadic behaviour makes the species vulnerable due to an increased exposure to predators and interaction with vehicles. This behaviour also brings humans and wildlife together and turtles are widely regarded as a positive connection with nature and eastern long-necked turtles are common pets throughout Australia. Hence, we used the eastern long-necked turtle as a model species in this study to assess the impacts of wetland loss in urban areas.

The aim of this study was to create a wetland inventory for Western Sydney, Australia and quantify changes in wetland abundance and surface area from 2010 – 2017. We quantified changes in primary land use (natural, urban or agricultural) over the same time period to determine if wetland loss is associated with increased urbanisation. Lastly, we then created species distribution models for the eastern long-necked turtle to determine changes in distribution throughout western Sydney over the last 20 years.

## Methods

### Study Area

This study focused on 10 Local Government Areas (LGAs) to produce a wetland inventory for western Sydney (NSW, Australia) (*Figure 1*). These LGA’s are currently undergoing significant urbanisation and land use change (NSW Planning & Statistics 2017). We classed land use into 3 categories; urban, natural and agricultural. Urban land was defined as developed areas with manmade infrastructure. Natural land was defined as areas with tree/shrub coverage or grassland. Agricultural land was defined as land used for farming, cropping and horticulture (Burgin *et al*. 2016).

**Figure 1:**
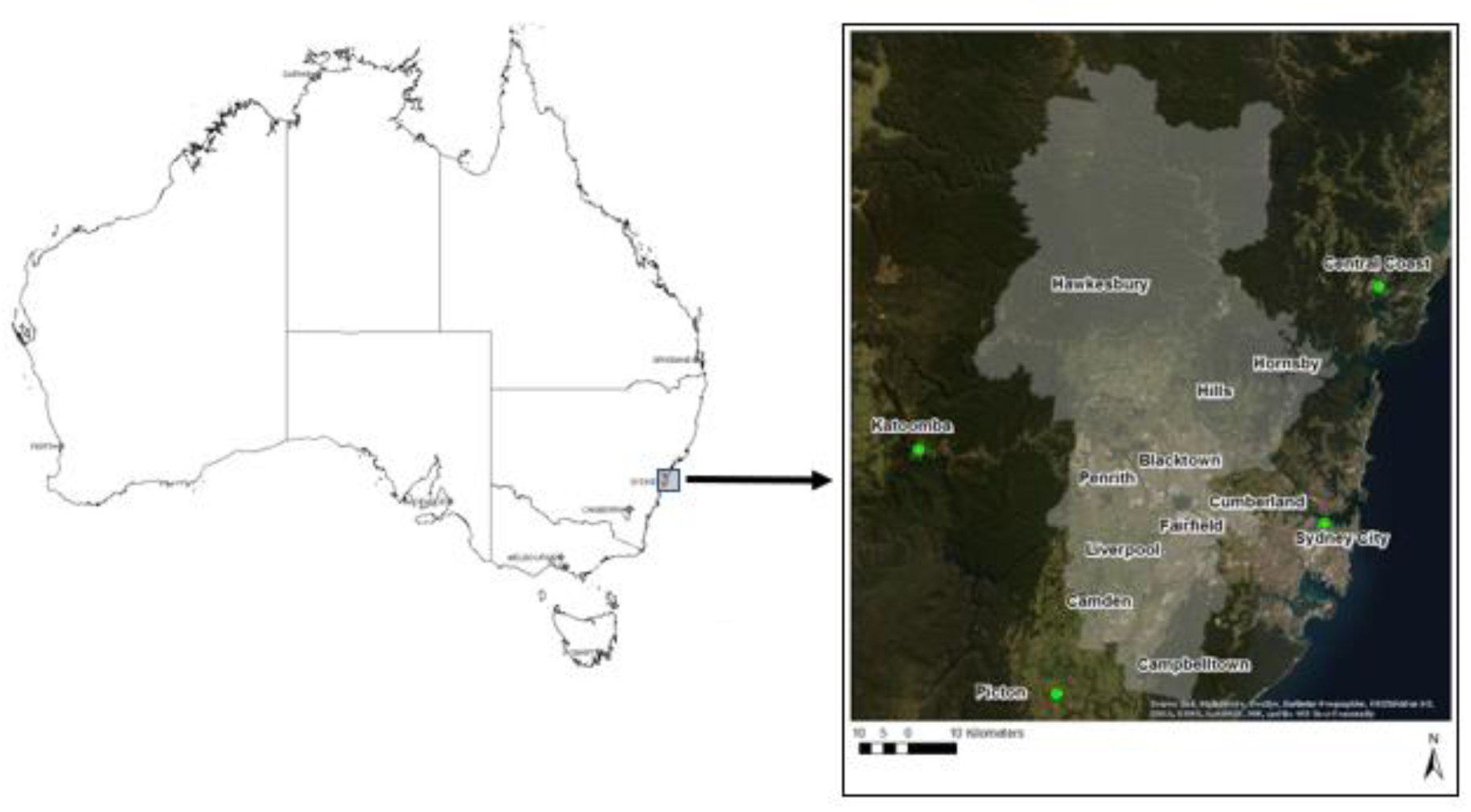
Map of western Sydney LGA’s sampled for this study (Google Imagery, 2019)

### Changes in Wetlands and Land Use

We used the online aerial imagery service NearMap (NearMap 2017) to record the number and size of every wetland in the above mentioned LGAs. A wetland was any form of water body visible such as a dam, pond and lake. Urban drains and rivers were not included, instead been noted as present. Each LGA was split up into suburbs, and grids with 50m zoom were established to methodically measure and record every wetland present (1,778 grids in total). Wetlands were measured with the area tool in NearMap, creating a total surface area in square meters of wetland for each suburb/LGA, this was then converted to a coverage percentage. Data was collected for the NearMap imagery available in April 2010 and then repeated in December 2017. Nearmap allows imagery from each time period to be changed, without changing the grid. Total square meters and percentage cover were then compared between 2010 and 2017 to observe changes. As well as counting wetlands, we recorded the land use of each grid as a cover percentage and compared between 2010 and 2017.

We created wetland hotspot maps using the Geographic Information System ZoaTrack (ZoaTrack 2017). Fixed kernel distribution (FK) (95% & 50%) and minimum convex polygon (MCP) (100% & 50%) analyses were used to create entire and core hotspots. Hotspot maps were created for both land use change and wetland loss data for each LGA using specific parameters; >10% loss in wetland coverage and >10% increase in urban land use cover. Wetland loss and land use change core hotspots were then overlayed to determine overlap.

### Species Distribution Modelling and Turtle Displacement

We used the Biodiversity and Climate Change Virtual Laboratory (BCCVL) to create species distribution models for the Eastern Long-Necked Turtle (ELNT) by associating known occurrences of the species and the environmental conditions of these areas. Occurrence only data (sightings) from across western Sydney was downloaded from the Atlas of Living Australia (ALA 2019). We filtered data to include only recent sightings (from the past 8 years), and split data into two periods; 2000-2007 (499 sightings) and 2010-2017 (844 sightings). The filtered data included all sexes and ages, but only live turtles. We chose four bioclimatic variables, downloaded from WorldClim v2.0 (Hijmans *et al*. 2005) at a spatial resolution of 30 arcsec (∼1 km). Variables relating to temperature and rainfall were chosen as these are most likely to affect movement and occurrences of the ELNT throughout wetlands in western Sydney. We used random forest (rf) algorithm models; a regression tree method with bootstrapping-based probability estimation, which allow for the presence-only data available for this study. This algorithm uses machine-learning method to create pseudo-absences and predict probability of environmental suitability of the species to an area. This algorithm merges a multitude of classification trees to produce highest consensus prediction (Rodrigues & Lima-Riberio 2018). Random forest is suitable for small datasets and is effective with few predictor variables (Cutler *et al*. 2007). We created a polygon within BCCVL for prediction, by providing our own constraints of western Sydney. This corresponds with the same area analysed for wetland loss and land use change. Random forest produced a geographic prediction of the probability of occurrence.

A turtle density estimate per square metre of wetland surface area was averaged from past research of eastern long-necked turtle population data (Ryan 2014; Ryan *et al*. 2015; Lovich *et al*. 2018). This resulted in a crude density estimate of 0.04 turtle per square metre of wetland surface area. Based on the loss of wetland surface area, an estimate of turtles affected was calculated for each LGA. Overall, presenting an estimate for the potential number of eastern long-necked turtles either displaced or deceased due to the loss and degradation of wetlands in western Sydney from 2010-2017.

## Results

### Changes in Wetland Density and Surface Area

Wetland surface area was largest in the LGA’s of Penrith, Hawkesbury and Cumberland between 2010 and 2017. Campbelltown, Fairfield and Hornsby had the smallest wetland surface area, all under 1,000,000 m^2^ for both 2010 and 2017. Excluding Campbelltown, Hawkesbury and Penrith, other LGA’s experienced a decrease in wetland surface area between 2010 and 2017. These decreases ranged from <1% (Cumberland) up to 21% (Blacktown), whilst the increases ranged from <1% (Hawkesbury) to 18.1% (Penrith) (*Figure 2*).

**Figure 2:**
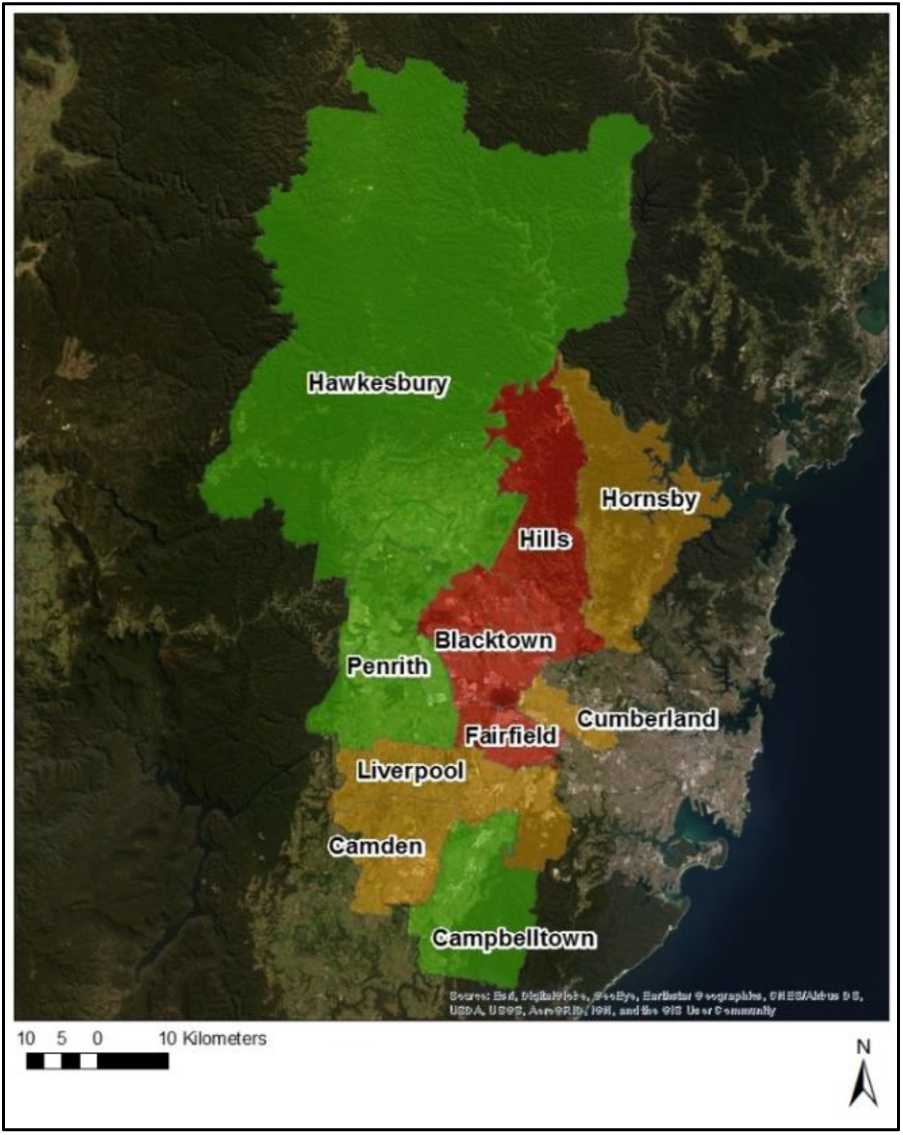
Change in western Sydney wetland surface area between 2010 & 2017 per LGA (>1% increase [green]; 0-10% decrease [orange]; >10% decrease [red])

Excluding Campbelltown and Cumberland, all LGA’s experienced a decrease in wetland density between 2010 and 2017. Cumberland experienced no change, whilst Campbelltown had an increase in wetland density (>5%). The majority of LGA’s had up to 10% loss of wetland density. Blacktown showed the largest change with a 25% decrease in the number of wetlands between 2010 and 2017 (*Figure 3*).

**Figure 3:**
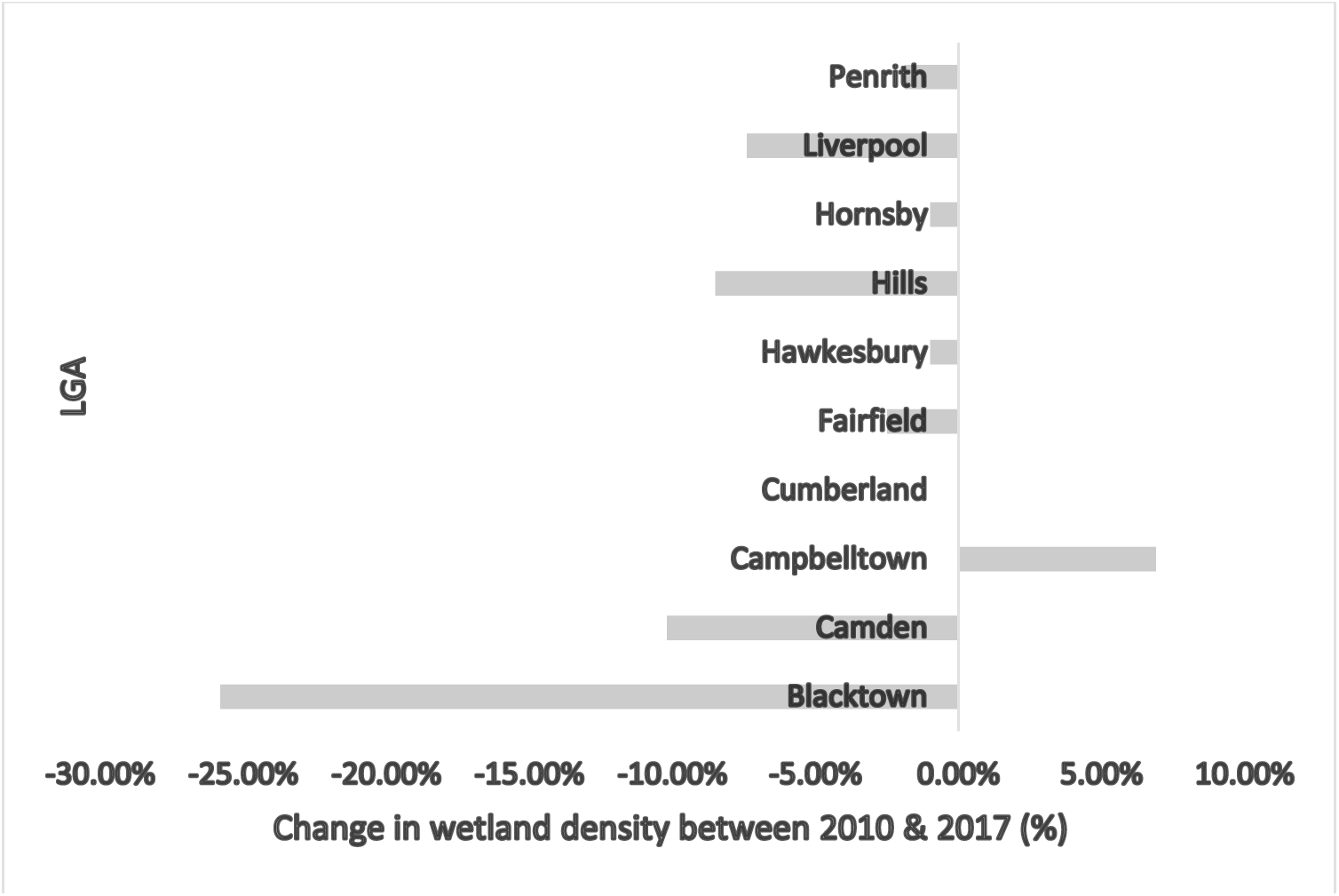
Change in wetland density (%) between 2010 & 2017 per LGA

### Changes in Land Use

Excluding Hornsby, all LGA’s experienced an increase in urban land use and a decrease in natural and agricultural land use. Hornsby had no change in land use category (*Figure 4*). Camden showed the largest increase in urban land use (12%). Blacktown, The Hills and Liverpool all gained over 5% of urban land. Camden lost the largest extent of natural land (11%). Blacktown, Liverpool and Penrith lost between 3% and 6% (*Figure 4*).

**Figure 4:**
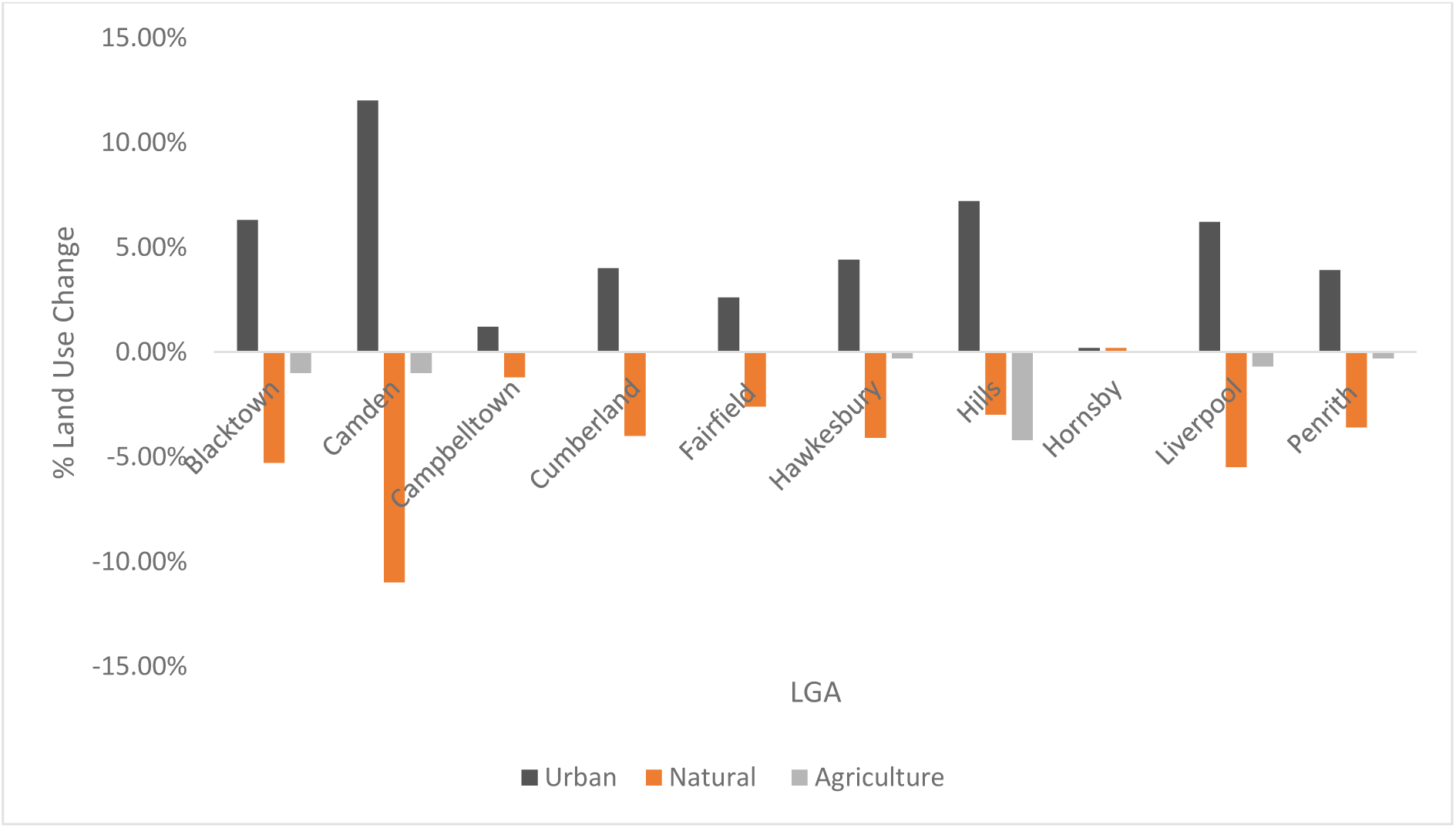
Land use change (%) between 2010 & 2017 per LGA

There is a significant overlap between the location of wetland loss and increased urbanisation (*Figure 5*). Blacktown, Penrith, The Hills and Liverpool are LGA’s that have suburbs with high rates of urbanisation and wetland loss. Fairfield has lost wetland surface area, but this was not associated with urbanisation. Camden shows a hotspot for urbanisation but not major reduction in wetland area from 2010-2017.

**Figure 5:**
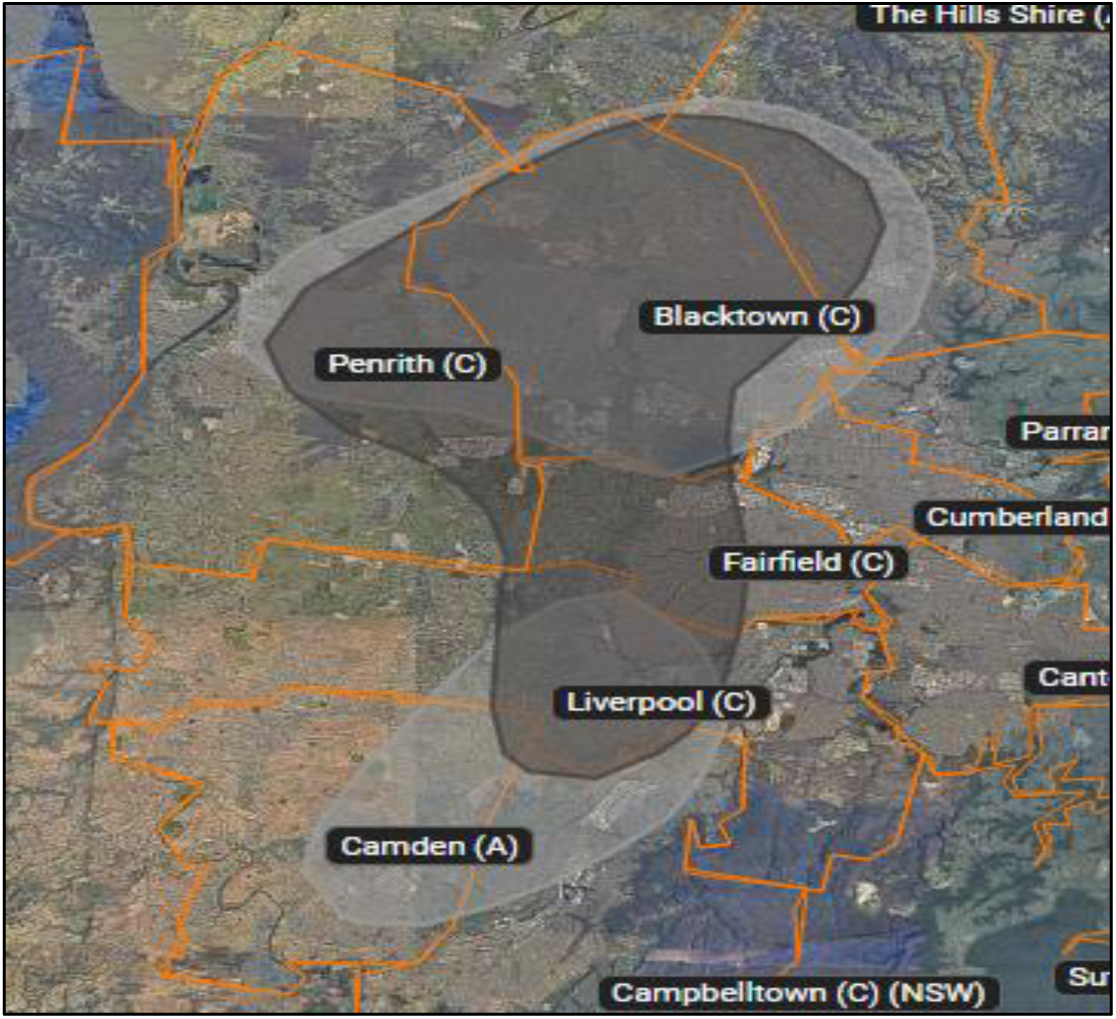
Overlap between wetland loss (dark grey) and urbanisation (light grey) hotspots (50% fixed kernel utilisation distribution) (Base Imagery: NearMap, 2018)

### Species Distribution Modelling

The species distribution models show a change in distribution of the eastern long-necked turtle throughout western Sydney between 2000-2017 (*Figure 6*). The distribution in 2000-2007 is less fragmented than 2010-2017. It is more condensed in the northern section of the study area in distinct hotspot patches in LGA’s such as Penrith, Hawkesbury, Blacktown and Liverpool. By 2010-2017 these hotspots move south-west with the species predicted to have isolated population groups as far south as Camden and Campbelltown LGA’s. The hotspots are lower in quantity and surrounded by areas with a very low prediction of species occurrence. The pattern of distribution correlates with the wetland loss hotspot suggesting a relationship (*Figure 5*).

**Figure 6:**
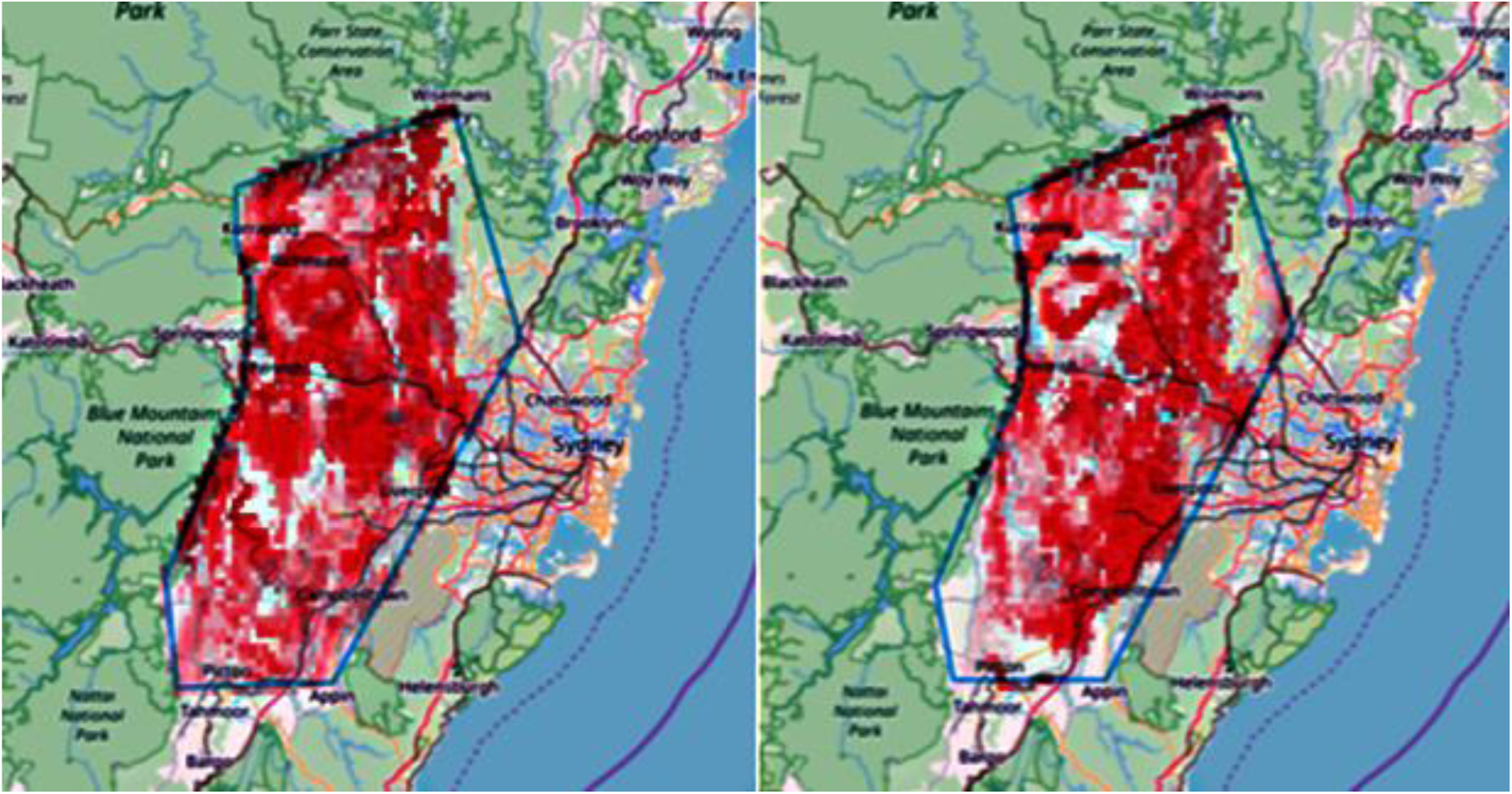
Species distribution models (rf) for the eastern long-necked turtle (*Chelodina longicollis*) for 2000-2007 (499 occurrences) & 2010-2017 (844 occurrences)

### Species Density Estimates

Based on decreases in wetland surface area, up to 15,000 turtles in Blacktown may have been displaced over the eight years, closely followed by Camden and the Hills with around 10,000 turtles each. Overall it is estimated that 40,000 individual eastern long-necked turtles were displaced or deceased due to wetland loss or degradation in western Sydney between 2010-2017. Some areas increased wetland surface area, and while this may provide habitat for turtles, these areas were also associated with a reduction in distribution (*Figure 6*), thus the quality of these constructed wetlands in these areas may not support long-term turtle populations.

## Discussion

Broad changes in both the number of wetlands and wetland surface area are a significant lost opportunity to integrate green infrastructure, green space and planned biodiversity conservation into the urban environment. In some areas of western Sydney, wetland loss has occurred rapidly with up to 21% of wetland surface area and 25% of wetland density decreasing over the eight-year period (*Figure 2; Figure 3*). Urban land use type increased on average, by 4.8% per LGA (*Figure 4*), and the hotspots for increased urbanisation almost directly matched the critical hotspots for wetland loss (*Figure 5*). Thus, development due to urban sprawl is the cause of wetland loss in western Sydney (Davis & Froend 1999; Burgin *et al*. 2016). Overall (excluding Penrith) ∼4.3% of wetland surface area and ∼5.2% of wetland density was lost in western Sydney between 2010-2017 (*Figure 2*; Figure 3), but the increases in wetland surface area in some LGAs (e.g. Penrith) was largely associated with development at the Penrith Lakes Scheme (mining) and flood and stormwater mitigation ponds (Penrith Lakes 2018). Both areas have effectively increased water storage, but it is unlikely that both areas serve as functional wetlands with complex ecosystems. We see some reduction in the distribution of turtles through that area (*Figure 6*) despite increases in water surface area (*Figure 6*). However, the long-term value of Penrith Lakes is significant. The 2000-hectare area was originally Cumberland woodland vegetation community but has functioned as a quarry from 1880 to 2013 (Penrith Lakes 2018). Since 2013, PLS has aimed to restore terrestrial and aquatic habitat to create an ecosystem that is both a replica of the original landscape with biodiversity corridors along the Nepean River, as well as provide green spaces for recreational purposes (Penrith Lakes 2018). In contrast, new developments appear to have paid little regard to integrating functional wetlands into the urban environment. Jordan Springs in Penrith LGA is an example where the potential of wetlands as green infrastructure could have been an important consideration prior to development. Jordan Springs underwent rapid landscape change between 2010 and 2017 and experienced an increase of 59,619.45m^2^ of wetland (+386.4%) but also 45% increase in urban land use (*Figure 4*). Land that was once natural bushland/grassland was converted into dense housing development. Originally there were no permanent wetlands present, however, two man-made wetlands were constructed with concrete floors and designed for stormwater retention and aesthetic/recreational purposes (Lendlease 2018). The largest wetland is the “Village Centre Lake” situated in the middle of the town centre and surrounded by a concrete pathway for cycling and homogenisation of reed species or bare rock patches around the circumference (*Figure 7*). Its value as complex green infrastructure is minimal.

**Figure 7:**
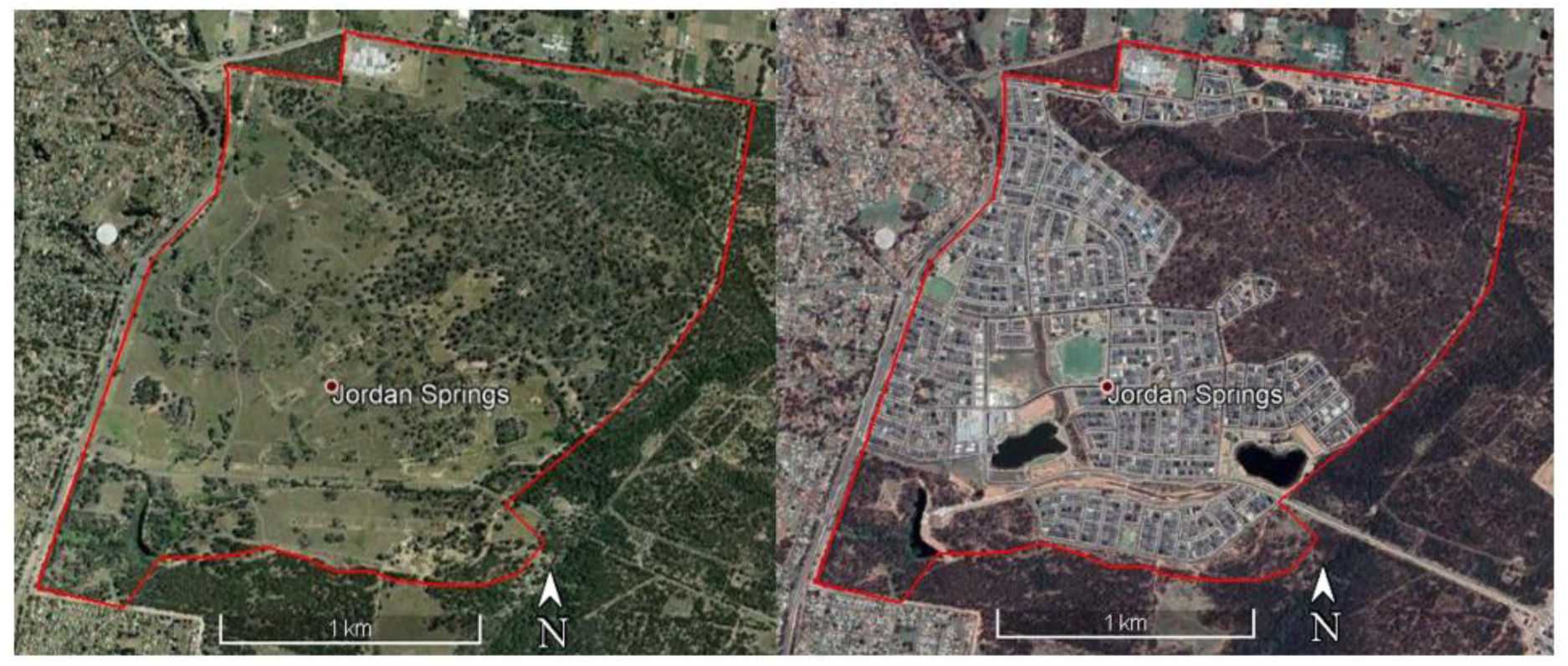
Jordan Springs 2010 (no wetlands) vs Jordan Springs 2017 (2 wetlands [+386.4%]) (Base Imagery: Google Earth 2020)

Constructed wetlands technology is an established green multi-purpose option for water management and wastewater treatment, with numerous effectively proven applications around the world and multiple environmental and economic advantages (Stefanakis 2019). These systems can function as water treatment plants, habitat creation sites, urban wildlife refuges, recreational or educational facilities, landscape engineering and ecological art areas (Stefanakis 2019). Construction of artificial functional wetlands is not new. For example, Marion council in Adelaide, Australia, developed a major stormwater use wetland in 2013, where a former Learner Driver Development Centre site at Oaklands Park was redeveloped into a functional wetland (Water Sensitive SA 2018). The wetland enables up to 200 million litres of stormwater to be used for irrigation each year at up to 30 council reserves and includes a range of habitats and depths and the site is now a haven for native birds, frogs, turtles and fish, as well as, an extensive range of aquatic and terrestrial invertebrates (Water Sensitive SA 2018). More than 85,000 aquatic and riparian zone plants have established since 2013 (Flinders University 2018). Integrated natural wetlands and constructed urban wetlands offer significant opportunities for green infrastructure and potential habitats for wildlife conservation that can help mitigate the negative effects of human activities on biodiversity decline. People experience biodiversity primarily where they live (Nilon et al. 2017). Urban planning and policy influences how people and communities experience and understand biodiversity.

With the increase of Citizen Science programs for species conservation, the capacity for the community to support conservation programs in urban areas is real and significant (Santori et al. 2020). Citizen Science also increases community engagement and education in the city and beyond (Santori et al. 2020). Daily interaction with nature engages people in conservation (Fuller and Irvine 2010) and has positive effects on physical and psychological health, social cohesion, crime reduction, environmental awareness, economic gain, and sense of belonging (Giles-Corti et al. 2005, Barton and Pretty 2010, Nilon et al 2017). Due to their long lifespan, slow movement, sturdiness, and wrinkled appearance, turtles are an emblem of longevity and stability in many cultures around the world (Ball 2002, Cirlot and Sage 2004). Urbanisation has long been at odds with wildlife, however ELNTs do better in suburban wetlands than in nature reserves (Roe and Georges 2007). Yet wetland removal is habitat destruction for ELNTs, and we estimated that up to 40,000 turtles were either killed or displaced over the period of this study. The ELNT is representative of the future of many common wetland dependent species; the population distribution changes are indicative of the way biodiversity is affected by wetland loss due to urbanisation (Figure 6). This species relies on both aquatic and terrestrial habitat at different stages of their life cycle, so the loss of wetlands and connectivity between them due to an increase in urbanisation can have severe effects on the population distribution (Rodrigues & Lima-Ribeiro 2018). Eastern Long Neck Turtle populations have declined by up to 91% in south-eastern Australia (Chessman 2011), however it does not have either a state or federal high-level conservation status. It inhabits all types of wetlands in western Sydney (Hull 2016), although is not considered in Impact Statements when development plans require the dewatering of a wetland. Community conservation groups, like Turtle Rescue NSW and Turtles Australia, are all that stands between complete extinction of common species due to habitat destruction during urban development (Figure 8). Western Sydney is still urbanising, this number will only increase as urban sprawl continues to move south and west (NSW Department of Planning & Statistics 2018). The occurrence probability patches in Camden and Campbelltown are ELNT hotspots and areas that should be prioritised for conservation and planning (Figure 6).

**Figure. 8.**
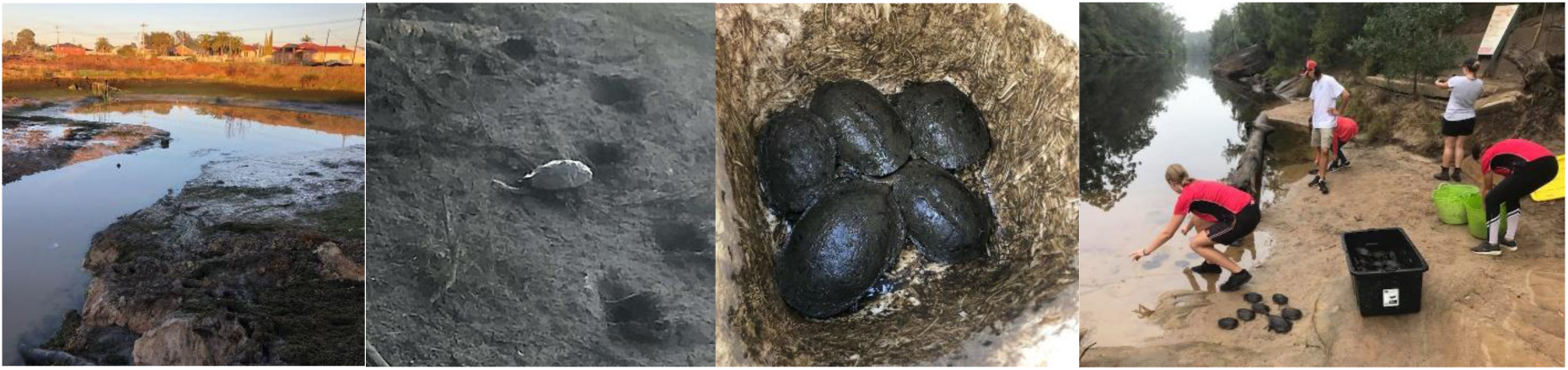
A wetland being dewatered in Western Sydney in Jun 2020. ELNTs emerge from wetlands as they are dewatered and must find other wetlands and are at risk of death from cars or dehydration in urban areas. Community conservation groups, like Turtle Rescues NSW, go in and collect turtles and release them in nearby creeks and rivers. (Photo credit: Turtle Rescues NSW)

The legislative requirements for developers removing wetlands varies throughout the world. Albert, Canada for example, implement a strategy where landowners must apply for a permit before undertaking any task that may negatively impact a wetland. However, as no wetland inventory exists, any illegal draining of wetlands cannot be monitored and mitigated, and between 1999-2009 80% of wetland loss was due to illegal draining (Clare & Creed 2014). Wetland inventory is crucial as a resource of accountability to enforce degradation and removal legislation, but also as a tool to assess the effectiveness of wetland policies. Western Sydney development is governed by the Biodiversity Conservation Act of 2016 (BC Act 2016). This act guides the NSW Wetlands Policy which has a list of 12 principles utilised for determining a “critical” wetland (DECCW 2010). Responsibility generally falls to the landowner to maintain wetlands; however, this policy does not consider the flow of water between wetlands and how wetlands are connected, or the potential of wetlands as biodiversity habitat. Developers must complete specific environmental assessments to gain approval from the local council, which interpret mitigative principles differently. These assessments generally focus only on the presence of threatened flora, fauna or communities which may be protected or potentially offset by wetlands of equivalent “values, functions and services”. If no offsets are available, then a price will be calculated via the Biodiversity Assessment Method (BAM) and this will be paid into the Biodiversity Conservation Trust as funding for conservation (DECCW 2010; BCT 2018). Constructed dams are not covered by this legislation regardless of age and are the most common form of wetland destroyed (Hansson et al. 2005; Hull 2016).

There is the necessity for functional wetlands that will meet the requirements of water treatment, sanitation and flood mitigation, and at the same time, provide green infrastructure in an urban environment (Stefanakis 2019). Nature-based solutions and, specifically, eco-friendly technology of natural and constructed wetlands are an ideal multi-purpose possibility that can maintain ecosystem services, provide habitat and resources for critical biodiversity, and enhance water circularity in the urban landscape (Masi, Rizzo & Regelsberger 2018). The integration of sustainable green technology and wetland inventory into effective government policy, which promotes responsibility and ownership of wetland nutrients, will support a balance between economic and biodiversity growth in an urbanising environment. Combined with “net biodiversity positive” land management, it places an economic value on green infrastructure like wetlands and the biodiversity associated with it. Wetlands also contribute to the well-being of the community by acting as urban green spaces, providing aesthetic appeal, landscape diversity and recreational opportunities. They can also contribute to cultural heritage, spiritual values and day-to-day living of Aboriginal and Torres Strait Islander peoples in Australia. Additionally, wetlands are accessible educational opportunities to learn about the environment, which promotes community engagement in conservation and supporting native biodiversity and sustainability.

## Acknowledgements

We thank Kane Durrant and Shane Davies from Turtle Rescues NSW. We thank Julia Ryeland and Kristen Petrov for technical aspect of this study. Financial support was provided by the Australian Research Council Linkage Grant Program (LP150100007), North-Central Catchment Management Authority, Yorta Yorta Aboriginal Corporation, Foundation for National Parks and Wildlife, Victorian Department of Land, Environment, Water and Planning, Winton Wetlands, Save Lake Bonney Group Inc.

